# Cognitive decline in aging parasitoid wasps

**DOI:** 10.64898/2026.02.03.703543

**Authors:** Hossein Kishani Farahani, Mathilde Lacombrade, Gabriel Madirolas Pérez, Coline Monchanin, Mathieu Lihoreau

## Abstract

Aging induces cognitive decline in humans and some other animals. For species that rely on learning and memory for reproduction, impaired cognitive functions may incur severe fitness costs. Here we report age-related cognitive decline in a solitary parasitoid wasp, *Venturia canescens*, that uses olfactory memories for host seeking and selection. We trained individual wasps to associate an odour with an oviposition reward, and compared their learning and memory performances at different stages of the reproductive life. Wasps between 6- and 14-day-old showed consistently poorer learning and reduced memory retention than young conspecifics, and this tendency increased with age. In this parasitoid insect, aging induces a precocious cognitive decline in reproductive females, which could severely impact their fitness through altered abilities to identify high quality hosts.

## Introduction

Aging induces gradual decline of physiological functions essential for survival and reproduction (Gilbert 2000). At the brain level, this often results in impaired cognitive functions (Martin 2011). Such cognitive decline is well-documented in humans, where older adults show reduced working memory, processing speed, and learning capacity (Zelinski and Burnight 1997), due to cellular alterations such as reduced dendritic branching and spine density (Duan et al. 2003), lower numbers of synapses (de Groot and Bierman 1987) or impaired synapse function (Barnes 1994). These consequences of aging are not specific to humans and can be observed across a wide range of species, including small-brained insects in which explorations of the links between well-defined cognitive traits and brain functions are facilitated (Promislow et al. 2022). Old drosophila flies show reduced learning and memory performances (Tamura et al. 2003) associated with a loss of neuronal synapses and a decreased number of fibers in the mushroom bodies of the brain (Liao et al. 2017), as well as changes in neurotransmitter release (Zhan et al. 2007).

While most attention has been dedicated to the mechanisms by which aging degrades learning and memory (Duan et al. 2003; Liao et al. 2017), relatively little is known about the fitness consequences of such decline. In principle aging can incur severe fitness costs when animals can reproduce throughout their lives, for instance through reduced parental care. These negative effects may be particularly important in solitary species where single adults provide all the parental care for their offspring. This is the case of many parasitoid insects that rely on learning and memory to choose a suitable host for their offspring development (Hoedjes et al. 2011). In these species, even a slight impairment of cognitive functions used to locate high quality hosts may importantly reduce the fitness of adults and their larvae.

Here we tested the effect of aging on the cognitive performances of a solitary parasitoid wasp *Venturia canescens*. Females of this species extensively rely on odour learning and memory to locate hosts and optimize their reproductive success by associating odours with best quality hosts (Hoedjes et al. 2011). We conditioned individual female wasps at various stages of their reproductive period (between 1- and 14-day-old) to associate an odor to a host. We then compared their learning performance and memory retention across ages.

## Material and methods

We ran the experiments in 2022-2023 at the University of Tehran, Iran. The *V. canescens* wasps were sourced in a laboratory culture originating from wild-caught individuals sampled in 2017 at Saveh (Markazi province, Iran). We raised the wasps and their hosts, the flour moth *Ephestia kuehniella*, at 25^°^C with a 16:8 (light: dark) photocycle under 50 ± 5% relative humidity. We obtained *E. kuehniella* eggs from the Insectary and Quarantine Facility of the Iranian Research Institute of Plant Protection and reared the larvae on a standard diet (48.5g wheat flour, 3g brewer yeast; (Kishani Farahani et al. 2019)). To obtain experimental subjects, we placed groups of 30 1-day-old *V. canescens* female wasps with ca. 200 5^th^ instar host larvae in a plastic box (30×20×20 cm) for 24h during which the wasps could lay eggs through telytokous parthenogenesis. We then kept the parasitized hosts in smaller boxes (5×5×3 cm) until the emergence of adult female wasps ca. four weeks later. Upon emergence, we feed fed these wasp 10% honey water solution (v/v).

We ran the experiments in a flight tunnel (200 × 50 × 50 cm) made of transparent plastic (for more details see (Kishani Farahani et al. 2021)) in an experimental room lit by 2000 lux white LEDs (Pars Shahab Lamp Co., Iran) (Kishani Farahani et al. 2019). Air was propelled inside the tunnel by a fan located at the upwind end and extracted outside using a fume hood at the downwind end (wind speed of 70 cm.s^-1^). The end part of the tunnel was divided into two decision chambers by a glass separator wall. Each chamber contained an odourant stimulus presented on a filter paper attached to a glass pipette placed vertically on a stand. For all experiments, we used orange and vanilla odours (97% pure odours: Adonis Gol Darou Group, Iran) (Kishani Farahani et al. 2021), for which the wasps had no innate preference (see Fig. S1).

### Conditioning

We measured the effect of aging on learning using olfactory conditioning assays in which the wasps had to associate an odour (orange) to a reward (oviposition) (Fig. 1). Prior to conditioning, wasps (N = 840) were individually exposed to a group of 15 host larvae (5^th^ instar) in cylindrical vials (2 cm ×10 cm) during 15 min (Fig. 1a). We used this early host exposure to make sure the wasps had some oviposition experience before the tests (Kishani Farahani et al. 2019). We then transferred 60 of these wasps into individual conditioning tanks (25 cm × 25 cm × 25 cm) with 30 5^th^ instar host larvae. Orange odour (conditioned stimulus, CS+) was pumped into the tanks at an air speed of 1 m.s^-1^. The wasps were maintained in these conditions for 2h during which they could associate the orange odour to host larvae (Fig. 1b). As negative control, a group of females was exposed to host larvae without orange odour and thus did not experience odour conditioning (unconditioned wasps, N = 420). We repeated this procedure for wasps aged between 1 to 14 day-old, an age range encompassing the first half of the typical lifespan of *V. canescens* adults and during which females continuously lay eggs (Harvey et al. 2001).

**Fig. 1:**
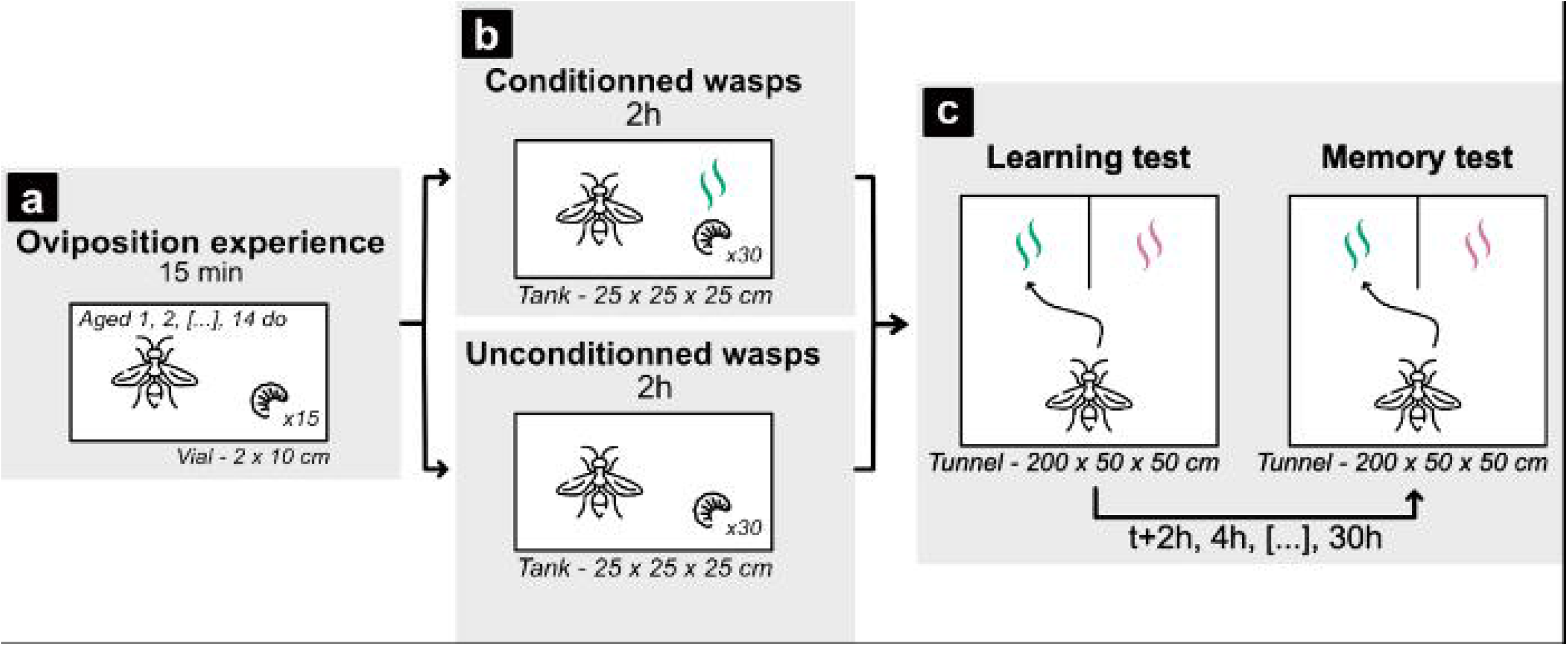
Experimental design. a) We placed wasps of different ages (60 per age class, N total = 840 individuals) with host larvae for 15 min to provide them an oviposition experience. b) We exposed a subset of these wasps (conditioned wasps, N = 420) to the orange odour (CS+) and host larvae (reward). The other subset of wasps (unconditioned wasps, N = 420) never experienced this association. c) We tested the learning performances of conditioned and unconditioned wasps by letting them choose between the orange odour (CS+) and a new odour (NOd, vanilla). To test for memory retention, we exposed the wasps to the same choice at different intervals after the first trial (by steps of 2h).

### Acquisition test

We assessed learning 15 min after conditioning by presenting wasps the conditioned odour (CS+: orange) and a new odour (NOd: vanilla) in the chambers of the flight tunnel (Fig. 1b). Every wasp that spent more than 3 consecutive minutes within 3 cm of the CS+ was considered as making a ‘correct choice’ and classified as learner (Kishani Farahani et al. 2021). Wasps that spent more than 3 minutes close to the NOd made an ‘incorrect choice’ and were classified as non-learners. Wasps that remain inactive or made no decision within 5 minutes of being introduced to the arena made ‘no choice’. We tested 420 conditioned and 420 unconditioned wasps across 14 age classes (Fig. 2a). We hypothesized that only conditioned wasps could make a correct choice based on the learned association between the host and the orange odour, and that learning performance would decrease with age.

**Fig. 2:**
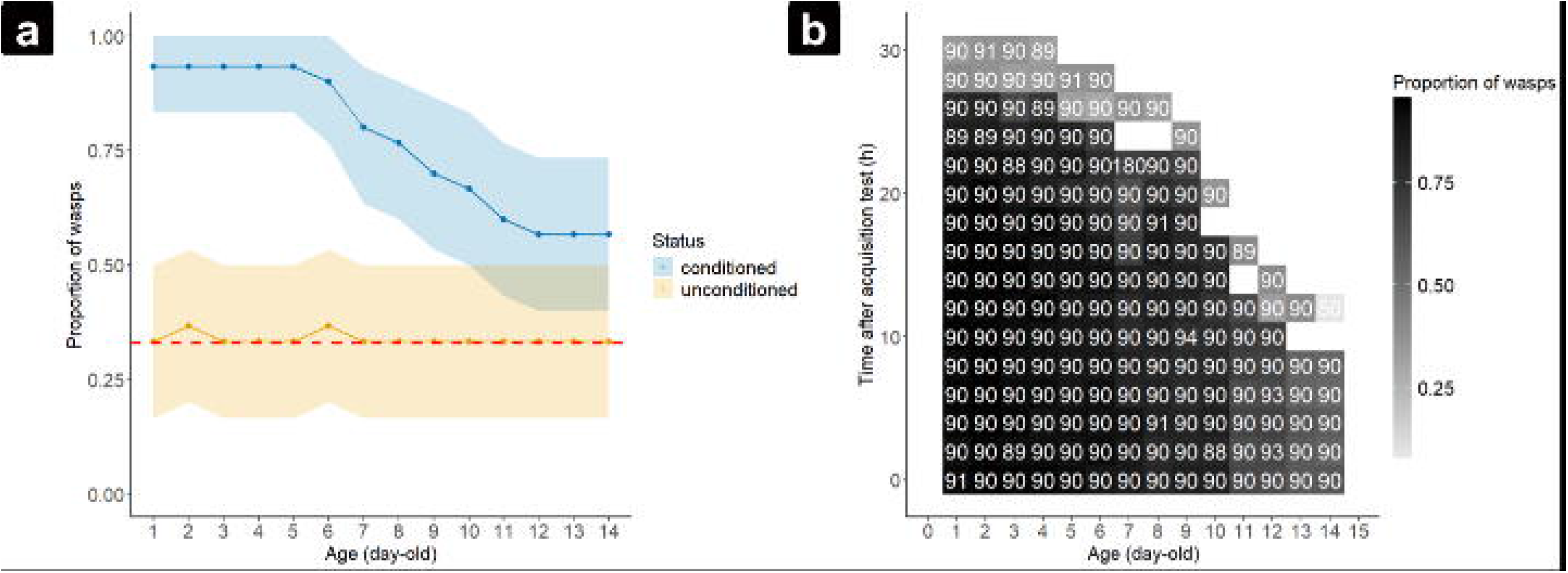
Results. a) Effect of age on learning. Proportion of wasps choosing the correct odour when conditioned (N = 420) or unconditioned (N = 420) across 14 age classes. Shaded areas represent 95% confidence intervals. The red dashed line represents chance level. b) Effect of age on memory retention. The heat map shows the proportion of wasps that made a correct choice in relation to age and the time interval between conditioning and memory test. The white characters indicate the sample size.

### Memory retention

We only tested the wasps that made a correct choice during the acquisition assay. We assessed memory retention at 14 age classes by observing the responses of the conditioned wasps either 0h, 2h, 4h, 6h, 8h, 10h, 12h, 14h, 16h, 18h, 20h, 22h, 24h, 26h, 28h, or 30h after conditioning (Fig. 1c). The responses of the wasps to the CS+ and the NOd were recorded in the flight tunnel using the same approach as above. Although we aimed to test 3 batches of 30 wasps of each age at each of the 15 retention times, conditioned responses were no longer detectable beyond specific post-conditioning intervals as wasps aged, which explains the lower sample sizes for higher age classes (Fig. 2b). Like before, we recorded whether the wasps made a correct choice, an incorrect choice, or no choice. We hypothesized that memory retention would decrease with age and time between conditioning and testing.

### Statistical analysis

We analyzed the data with R 4.3.3. From the raw data (Dataset S1 available upon request), we extracted the first choice of each wasp (CS+: 1, NOd: 0) in the acquisition and memory retention tests. We evaluated the influence of age (1-to 14 day-old) and conditioning status (conditioned or unconditioned) on acquisition using a generalized linear model (GLM). We then assessed the effect of age and the delay between tests (2 to 30 h after learning test) on memory retention using a GLM. Both GLMs had a binomial error structure (R package lme4) and were followed by an ANOVA.

## Results

### Learning performance decreased with age

We first assessed the effect of aging on learning by testing wasps at 14 age classes (Fig. 2a). In the conditioned wasps, the proportion of correct choices decreased with age (GLM, age × conditioning: Chi^2^=31.98 df=13, p=0.002). The proportion of correct choices was initially high (average of 93.33%; CI: 83.33-100%) in 1-to 6-day-old wasps, indicating successful conditioning. However, it progressively dropped with age to reach a minimum of 56.67% (CI: 40-73.33%) in 12-to 14-day-old wasps. As expected, the proportion of correct choices in unconditioned wasps remained at chance level (ca. 33.33%, red dashed line in Fig. 2a) for all age classes.

### Memory retention decreased with age and time post conditioning

We next measured the effect of aging on memory retention by testing conditioned wasps that successfully learned the odour-reward association (‘correct choice’ in acquisition test) at 16 different times post conditioning for each of the 14 age classes (Fig. 2b). The percentage of wasps that chose the correct odour decreased with age and with time post-conditioning (GLM, time between times × age: Chi^2^=256, df=142, p<0.001). Strikingly at 2h post-conditioning, 91.11% (CI: 84.44 - 96.66%) of the 1-day-old wasps made a correct choice whereas only 55.55% (CI: 45.55 – 65.55%) of the 14-day-old wasps did so. This effect of aging was even more pronounced for longer intervals post-conditioning. For instance, at 12h post-conditioning, 93.33% (CI: 87.78 –97.78%) of the 1-day-old wasps still made a correct choice whereas only 8% (CI: 2 – 16%) of 14-day-old wasps did so.

## Discussion

Older wasps had lower learning performances and shorter memory retention than younger ones. This decline of olfactory cognition during the first half of the typical reproductive period for this species suggests important fitness costs through reduced abilities to identify high quality hosts.

The learning rate of the parasitoid wasps dropped from 93.33% in 1-day-old wasps to 56.67% in 14-day-old ones. Impaired learning has already been reported in aged insects (Brown and Strausfeld 2009; Iliadi and Boulianne 2010), including other social hymenopteras such as wasps (Mattens et al. 2025) and honey bees (Behrends et al. 2007). However, in these social insects, impaired cognition has been associated to division of labour that characterizes social organization rather than chronological age per se like in the present study, since learning abilities decline faster in foragers than in same-aged nurses (Behrends et al. 2007) and foragers can recover learning performances after reverting into nursing tasks.

Aging in our parasitoid wasps also reduced memory retention by 36% only 2h after conditioning, and up to 85% 12h after conditioning. This dramatic decay of memory retention in old wasps is much stronger than previous reports in social wasps and bees (Behrends et al. 2007). This could be due to differences in experimental approaches. While in our experiment, we conditioned parasitoid wasps by exposing them to the host only once (2h), in previous studies the wasps were conditioned through repeated exposures (Mattens et al. 2025), a procedure that can promote memory consolidation (Behrends et al. 2007).

Our observations thus add parasitoid wasps to the list of species demonstrating learning and memory impairments with aging. In *V. canescens*, aging may incur important fitness costs since reproductive females can manifest reduced cognitive performances as soon as 7-day-old. Indeed, for parasitic wasps, host finding is directly linked to the number of offspring they can produce through the possibility to maximize the production of progeny per unit of time (Godfray 1994). Accordingly, females conditioned to odours associated to hosts tend to find faster and more reliably high quality hosts than unconditioned conspecifics (Papaj and Vet 1990). They also tend to parasitize more eggs and produce more offspring (Dukas and Duan 2000). Our findings in a non-model insect species highlights the prevalence of cognitive aging across the animal kingdom and emphasize the need to compare species with different life histories in order to clarify its fitness consequences. Simple organisms with rich cognitive repertoires and short generation times, such as insects, can greatly facilitate this exploration.

## Acknowledgments

We thank Mr. Poria Abroon, who helped with the maintenance of insect cultures and some of the behavioural trials. While writing MLa, GMP, CM and MLi were supported by the European Research Council (ERC Cog BEE-MOVE GA101002644 to ML).

## References

Barnes CA (1994) Normal aging: regionally specific changes in hippocampal synaptic transmission. Trends Neurosci 17:13–18. 10.1016/0166-2236(94)90029-9

Behrends A, Scheiner R, Baker N, Amdam GV (2007) Cognitive aging is linked to social role in honey bees (Apis mellifera). Exp Gerontol 42:1146–1153. 10.1016/j.exger.2007.09.003

Brown S, Strausfeld N (2009) The effect of age on a visual learning task in the American cockroach. Learn Mem 16:210–223. 10.1101/lm.1241909

de Groot D, Bierman E (1987) Numerical changes in rat hippocampal synapses. An effect of “ageing”? Acta Stereol 6:53–58

Duan H, Wearne SL, Rocher AB, et al (2003) Age-related dendritic and spine changes in corticocortically projecting neurons in macaque monkeys. Cereb Cortex 13:950–961. 10.1093/cercor/13.9.950

Dukas R, Duan JJ (2000) Potential fitness consequences of associative learning in a parasitoid wasp. Behav Ecol 11:536–543. 10.1093/beheco/11.5.536

Gilbert SF (2000) Aging: the biology of senescence. In: Developmental biology. 6th edition. Sinauer Associates

Godfray HCJ (1994) Parasitoids: behavioral and evolutionary ecology. Princeton University Press

Guarente L, Kenyon C (2000) Genetic pathways that regulate ageing in model organisms. Nature 408:255–262. 10.1038/35041700

Harvey JA, Harvey IF, Thompson DJ (2001) Lifetime reproductive success in the solitary endoparasitoid, Venturia canescens . J Insect Behav 14:573–593. 10.1023/A:1012219116341

Hoedjes KM, Kruidhof HM, Huigens ME, et al (2011) Natural variation in learning rate and memory dynamics in parasitoid wasps: opportunities for converging ecology and neuroscience. Proc R Soc B 278:889–897. 10.1098/rspb.2010.2199

Iliadi KG, Boulianne GL (2010) Age-related behavioral changes in drosophila. Ann N Y Acad Sci 1197:9–18. 10.1111/j.1749-6632.2009.05372.x

Kishani Farahani H, Moghadassi Y, Alford L, Van Baaren J (2019) Effect of interference and exploitative competition on associative learning by a parasitoid wasp: a mechanism for ideal free distribution? Anim Behav 151:157–163. 10.1016/j.anbehav.2019.03.017

Kishani Farahani H, Moghadassi Y, Pierre J-S, et al (2021) Poor adult nutrition impairs learning and memory in a parasitoid wasp. Sci Rep 11:16220. 10.1038/s41598-021-95664-6

Liao S, Broughton S, Nässel DR (2017) Behavioral senescence and aging-related changes in motor neurons and brain neuromodulator levels are ameliorated by lifespan-extending reproductive dormancy in drosophila. Front Cell Neurosci 11:. 10.3389/fncel.2017.00111

Martin GM (2011) The biology of aging: 1985–2010 and beyond. FASEB J 25:3756–3762. 10.1096/fj.11-1102.ufm

Mattens A, Christiaens H, Debeuckelaere K, et al (2025) Age-related differences in learning, memory and brain plasticity in workers of the common wasp Vespula vulgaris. J Exp Biol 229:jeb.251673. 10.1242/jeb.251673

Papaj DR, Vet LEM (1990) Odor learning and foraging success in the parasitoid,Leptopilina heterotoma. J Chem Ecol 16:3137–3150. 10.1007/BF00979616

Promislow DEL, Flatt T, Bonduriansky R (2022) The biology of aging in insects: from Drosophila to other insects and back. Annu Rev Entomol 67:83–103. 10.1146/annurev-ento-061621-064341

Tamura T, Chiang A-S, Ito N, et al (2003) Aging specifically impairs amnesiac-dependent memory in Drosophila. Neuron 40:1003–1011. 10.1016/S0896-6273(03)00732-3

Zelinski EM, Burnight KP (1997) Sixteen-year longitudinal and time lag changes in memory and cognition in older adults. Psychol Aging 12:503–513. 10.1037/0882-7974.12.3.503

Zhan M, Yamaza H, Sun Y, et al (2007) Temporal and spatial transcriptional profiles of aging in Drosophila melanogaster. Genome Res 17:1236–1243. 10.1101/gr.6216607

