## Supplementary Figure S1 for "Cognitive decline in aging parasitoid wasps"

**Supplementary Materials**

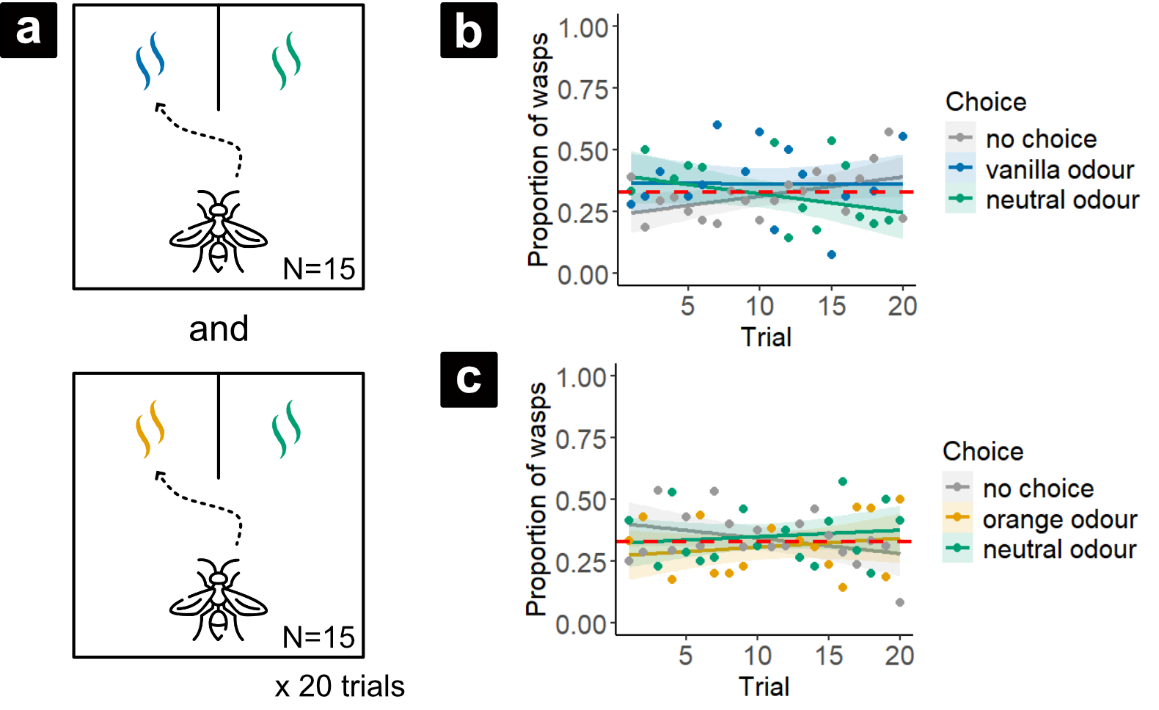

**Fig. S1**: **Test for innate odour preference.** a) Before the study, we controlled for an absence of innate attraction by the wasps to the odours used for conditioning (vanilla and orange). We gave individual wasps a choice between a filter paper scented with 1μl of either vanilla or orange odour in one chamber of the flight tunnel (referred to *vanilla* or *orange odour* and respectively colored in blue or orange in Fig. S1) and an unscented filter paper in the other chamber of the flight tunnel (referred to *neutral odour* and colored in green in Fig. S1). We placed the test wasp (N = 30) at the start zone of the tunnel and allowed it to choose between the two chambers for 15 minutes. We then repeated the procedure across 20 consecutive trials. Any wasp that spent more than 3 consecutive minutes within 3 cm around the scented filter paper (landed or hovering around) was considered as ‘making a choice’ (a wasp landing on an odour site for more than 3 minutes typically remains longer than 15 minutes) (Kishani Farahani et al. 2021). Any wasp that did not fly in the tunnel after 5 minutes was considered as ‘making no choice’ (referred as *no choice* and colored in grey in Fig. S1). All behavioural data were recorded by visual observation. b) We found no significant preference for vanilla or orange over the neutral odour and this did not change through trials (GLMM with *the trial and the presented odour* as fixed factors and *wasp_id* as random factor, *presented odor*: Chi² = 2.26, df = 1, p = 0.13; *trial*: Chi² = 0.39, df = 1, p = 0.53). The red dashed line represents the theoretical percentage in case of a random choice.
